# Effects of arm swing amplitude and lower limb asymmetry on motor variability patterns during treadmill gait

**DOI:** 10.1101/2021.09.24.461689

**Authors:** Christopher A. Bailey, Allen Hill, Ryan Graham, Julie Nantel

## Abstract

Motor variability is a fundamental feature of gait. Altered arm swing and lower limb asymmetry (LLA) may be contributing factors having been shown to affect the magnitude and dynamics of variability in spatiotemporal and trunk motion. However, the effects on lower limb joints remain unclear.

Full-body kinematics of 15 healthy young adults were recorded during treadmill walking using the Computer-Assisted Rehabilitation Environment system. Participants completed six trials, combining three arm swing (AS) amplitude (normal, active, held) and two LLA (symmetrical, asymmetrical) conditions. The mean standard deviation (meanSD), maximum Lyapunov exponent (λ_max_), detrended fluctuation analysis scaling exponent of range of motion (DFAα), and sample entropy (SaEn) were computed for tridimensional trunk, pelvis, and lower limb joint angles, and compared using repeated-measures ANOVAs.

Relative to normal AS, active AS increased meanSD of all joint angles, λ_max_ of frontal plane hip and ankle angles, and SaEn of sagittal plane ankle angles. Active AS, however, did not affect λ_max_ or SaEn of trunk or pelvis angles. LLA increased meanSD of sagittal plane joint angles, λ_max_ of Euclidean norm trunk angle and of lower limb joint angles, and SaEn of ankle dorsiflexion/ plantarflexion, but decreased SaEn of tridimensional trunk angles and hip rotation in the slower moving leg.

Alterations in lower limb variability with active AS and LLA suggest that young adults actively exploit their lower limb redundancies to maintain gait. This appears to preserve trunk stability and regularity during active AS but not during LLA.

## 1. Introduction

Motor variability in gait, the natural biological variability from stride to stride in motor outputs like movement time and kinematics (Newell and Slifkin, 1998), changes with older age (Beauchet et al., 2017; Buzzi et al., 2003; Kang and Dingwell, 2009; Kurz and Stergiou, 2003). Measurable features of motor variability beyond magnitude include dynamical stability (e.g. maximum Lyapunov exponent measuring local dynamic stability), persistence (e.g. scaling exponent from detrended fluctuation analysis), and regularity (e.g. sample entropy). For these features, older adults are reported to have lower local dynamic stability of the trunk and lower limb (Buzzi et al., 2003; Kang and Dingwell, 2009), no difference in persistence of fluctuations in stride length, time, or speed (Dingwell et al., 2017), and lower regularity of the knee and hip kinematics in the sagittal plane (Kurz and Stergiou, 2003). Furthermore, older adults who possess high motor variability have been shown to be more likely to fall (Callisaya et al., 2011; Hausdorff et al., 2001; Toebes et al., 2012), although some variability is beneficial since having too little can also leads to falls (Beauchet et al., 2009; Brach et al., 2005). These findings suggest that a loss of dynamic stability and regularity in kinematics could be a potential mechanism for incurring a fall during gait.

Prior studies indicate that older adults walk with smaller arm swing amplitude (Mirelman et al., 2015) and larger lower limb asymmetry (Aboutorabi et al., 2016) relative to young adults, meaning arm swing and lower limb asymmetry could be factors that contribute to motor variability. In young adults, actively increasing arm swing amplitude has been found to increase local dynamic stability of the trunk relative to normal arm swing (Hill and Nantel, 2019; Wu et al., 2016), increase the magnitude of variability in step time, length, and width (Hill and Nantel, 2019; Siragy et al., 2020), but not affect variability of hip knee and ankle angles (Wu et al., 2016). As discussed by Hill and Nantel (2019), active arm swing may stabilize the trunk by increasing angular momentum and its resistance to change or by increased attention to the movement of the torso and upper limbs. Loss of arm swing amplitude could then destabilize the trunk, however, studies of steady-state gait with restricted arm swing saw no such effect (Bruijn et al., 2010; Hill and Nantel, 2019). As for lower limb asymmetry, studies have shown in young adults that split-belt treadmill induced asymmetry decreases margin of stability (Buurke et al., 2018; Darter et al., 2018), decreases local dynamic stability of the trunk (Hill and Nantel, 2019), and increases magnitude of variability in step length (Hill and Nantel, 2019; Siragy et al., 2020). Since lower limb joint variations are likely responsible for spatiotemporal variability, it appears that exploitation of lower limb redundancy may not be sufficient to dynamically stabilize the trunk during asymmetric gait.

A key limitation presently in these investigations of arm swing amplitude and lower limb asymmetry is that motor variability has mainly been measured by spatiotemporal and trunk features and not across the full kinematic chain. To our knowledge, only Wu et al. (2016) investigated the influence of one of these factors on variability of lower limb joints in healthy young adults, finding no differences between active and normal arm swing. However, mean gait speed was relatively slow in their study (0.74-0.82 m/s) and increased with active arm swing, meaning gait speed influences on stride-to-stride variability (Dingwell and Marin, 2006) could have influenced their findings. Further exploration of the concurrent trunk, pelvis, and lower limb variability adjustments in young adults could help better understand how motor variability in gait emerges in older adults. Thus, we investigated in this study: 1) how arm swing amplitude and lower limb asymmetry alter variability of trunk, pelvis, and lower limb joint angles in gait and 2) if arm swing amplitude influences changes in variability of joint angles attributed to lower limb asymmetry. We hypothesized that variability of lower limb joint angles would not change with active arm swing, that magnitude of variability would increase and local dynamic stability would decrease with lower limb asymmetry, and that there would be no interactions between active arm swing and lower limb asymmetry.

## 2. Methods

### 2.1 Participants

Fifteen healthy young adults (8 males; 23.4 ± 2.8 years; 72.3 ± 13.5 kg; 1.70 ± 0.08 m) were recruited as a convenience sample from the Ottawa area as part of Hill and Nantel (2019). Participants were excluded if they had a musculoskeletal injury in the preceding six months, or any chronic neurological or orthopaedic disorders. Participants self-reported as right-hand dominant except for one participant who self-reported as ambidextrous. Each participants provided written informed consent, which followed the Declaration of Helsinki and was approved by the Ottawa Health Science Network Research Ethics Board (20170291-01H) and by the University of Ottawa Research Ethics Board (A06-17-03).

### 2.2 Procedure

Each participant completed gait trials in the Computer Assisted Rehabilitation Environment (CAREN) (CAREN-Extended, Motekforce Link, Amsterdam, NL). This combined a split-belt treadmill (TM-09-P-MOTEK, Motekforce Link, Amsterdam, NL) instrumented with a force plate (sampled at 1000 Hz; Bertec Corp., Columbus, OH) and a 12-camera optoelectronic motion capture system (sampled at 100 Hz; MX T20S, Vicon, Oxford, UK). Markers were positioned on the full body as described previously (Collins et al., 2009; Wilken et al., 2012). For each trial, the participant walked at 1.2 m/s for 200 seconds under one of three arm swing conditions (normal, held, active) and one of two symmetry conditions (symmetric, asymmetric). For held arm swing, the participant was instructed to “hold [their] arms still along [their] sides without shoulder tension and arm stiffness”. For active arm swing, the participant was instructed to “swing [their] arms forward to be horizontal at peak forward swing”. For asymmetrical gait, the participant walked at 0.96 m/s with their right leg while walking at 1.2 m/s with their left leg (0.8:1 ratio). Conditions were randomized and the participant completed one trial for each combination of arm swing and symmetry conditions.

### 2.3 Data analysis

Marker trajectories were low-pass filtered (10 Hz, Butterworth, zero-lag, 4^th^ order) and used to model tridimensional trunk (flexion, bending, rotation), pelvis (tilt, obliquity, rotation), hip (flexion, abduction, rotation), knee (flexion, valgus), and ankle (dorsiflexion, inversion) angles in Visual3D (C-Motion, Germantown, MD, USA) as previously described (Collins et al., 2009; Wilken et al., 2012). Trunk and pelvis angles were modeled relative to the global coordinate system. Ground reaction force data were low-pass filtered (20 Hz, Butterworth, zero-lag, 4^th^ order) and combined with selected kinematic features (foot position relative to pelvis, foot velocity relative to pelvis and the laboratory, foot acceleration, and knee angle) in a logistic classification model to identify heel strike events which defined the start and end of each stride; these events were manually inspected and corrected as needed.

Using MATLAB (R2020b, MathWorks Inc., Natick, MA, USA), the first 25 seconds of data in each trial were discarded to account for time needed for the participant to reach a steady-state, and joint angle variability outcomes (magnitude of variability, local dynamic stability, statistical persistence, regularity) were quantified for the subsequent 125 strides. For each degree of freedom, the magnitude of variability was quantified by the mean standard deviation (meanSD), calculated by normalizing continuous series to 101 points (0-100%) per stride, finding the standard deviation between strides at each normalized point, then taking the mean value. Local dynamic stability was quantified by the maximum short-term finite-time Lyapunov exponent (λ_max_) using the method of Rosenstein et al. (1993). Continuous series were normalized to 12500 points (100 per stride on average), then λ_max_ was computed with 5 embedding dimensions at a lag of 10 points from 0-0.5 strides (50 points) (Buzzi et al., 2003; Wu et al., 2016). λ_max_ measures the divergence of neighbouring trajectories with higher positive values indicative of higher divergence and lower local dynamic stability. Statistical persistence was quantified by the detrended fluctuation analysis scaling exponent (DFAα) (Dingwell et al., 2017), calculated from 125 consecutive joint angle range of motion values. DFAα is non-negative and unitless, with values > 0.5 indicative of persistence (a fluctuation is typically followed by a fluctuation in the same direction), values < 0.5 indicative of anti-persistence (a fluctuation is typically followed by a fluctuation in the opposite direction), and values ∼ 0.5 indicative of no correlation between fluctuations. Regularity was quantified by the sample entropy (SaEn), computed with 2 embedding dimensions and a 0.15 tolerance distance (Costa et al., 2003). SaEn can be investigated at several different scales using a multiscale function; we selected a factor of 4 which is believed to be approximately where entropy of physiological signals stabilizes during slow, normal, and fast walking speeds (Costa et al., 2003). SaEn is non-negative and unitless, with higher values indicative of lower regularity. In supplement, variability outcomes were also calculated from Euclidean norm angles for each joint and can be found in Supplementary Table 1.

### 2.4 Statistical analysis

Using SPSS (v27, IBM, Armonk, NY, USA), normality of joint angle variability outcomes was confirmed via Kolmogorov-Smirnoff tests. Repeated measures ANOVAs were conducted on each outcome to test for within-subjects effects of Swing (normal, active, held) and Symmetry (symmetric, asymmetric), as well as Swing^*^Symmetry interactions. For each ANOVA, sphericity was inspected and, when violated, Greenhouse-Geisser corrections were applied. We used the Benjamini-Hochberg procedure to control the false discovery rate due to multiple comparisons (Benjamini and Hochberg, 1995); at an *a priori* alpha of 0.050, the critical p-value was adjusted to 0.010 (240 p-values: 3 statistical effects ^*^ 20 individual joint angles ^*^ 4 variability outcomes). Partial eta squared effect sizes (η^2^) were computed and small, medium, and large effect sizes were defined with thresholds of η^2^ = 0.01, η^2^ = 0.06, and η^2^ = 0.14 respectively (Cohen, 1977). Pairwise post-hoc comparisons were made at a critical p-value of 0.010.

## 3. Results

### 3.1 meanSD, magnitude of variability (Table 1)

There were no significant Swing^*^Symmetry interactions on meanSD (p ≥ 0.010).

**Table 1.** Mean standard deviation (meanSD) of joint angles performed under six gait conditions varying by arm swing (normal, active, held) and symmetry (symmetric, asymmetric) Values are group means (inter-individual standard deviation). P-values of main and interaction effects are provided, with effect sizes (η^2^) for significant effects (p < 0.010).

| Joint | Degree of freedom | Symmetric |  |  | Asymmetric |  |  | Swing | Symmetry | Interaction |
| --- | --- | --- | --- | --- | --- | --- | --- | --- | --- | --- |
|  |  | Normal | Active | Held | Normal | Active | Held |  |  |  |
| Right hip | FE <sup>a*</sup> | 1.130 (0.212) | 1.880 (0.592) | 1.252 (0.310) | 1.550 (0.495) | 1.947 (0.601) | 1.571 (0.339) | .002, $\eta^2=0.484$ | <.001, $\eta^2=0.650$ | .085 |
| | AA <sup>a*</sup> | 0.814 (0.123) | 1.239 (0.320) | 0.826 (0.137) | 0.955 (0.218) | 1.269 (0.320) | 0.930 (0.150) | <.001, $\eta^2=0.645$ | .001, $\eta^2=0.529$ | .224 |
| | Rotation <sup>a</sup> | 1.100 (0.177) | 1.661 (0.468) | 1.130 (0.173) | 1.203 (0.178) | 1.617 (0.396) | 1.253 (0.248) | <.001, $\eta^2=0.688$ | .084 | .141 |
| Right knee | FE <sup>a*</sup> | 1.884 (0.423) | 2.773 (0.631) | 2.026 (0.498) | 2.565 (1.270) | 2.920 (0.711) | 2.457 (0.618) | .001, $\eta^2=0.392$ | <.001, $\eta^2=0.680$ | .246 |
| | VV <sup>a*</sup> | 0.661 (0.159) | 0.908 (0.264) | 0.655 (0.135) | 0.754 (0.212) | 0.900 (0.253) | 0.744 (0.188) | <.001, $\eta^2=0.580$ | <.001, $\eta^2=0.601$ | .260 |
| Right ankle | DfPf <sup>a*</sup> | 1.525 (0.329) | 2.084 (0.405) | 1.588 (0.278) | 1.817 (0.333) | 2.192 (0.377) | 1.800 (0.333) | <.001, $\eta^2=0.631$ | .003, $\eta^2=0.469$ | .247 |
| | InvEv <sup>a</sup> | 1.083 (0.241) | 1.456 (0.323) | 1.182 (0.424) | 1.180 (0.301) | 1.457 (0.314) | 1.127 (0.265) | <.001, $\eta^2=0.601$ | .676 | .186 |
| Left hip | FE <sup>a*</sup> | 1.142 (0.186) | 1.887 (0.706) | 1.249 (0.302) | 1.371 (0.241) | 1.983 (0.606) | 1.489 (0.342) | .002, $\eta^2=0.491$ | .008, $\eta^2=0.404$ | .362 |
| | AA <sup>a</sup> | 0.818 (0.126) | 1.239 (0.314) | 0.850 (0.150) | 0.930 (0.192) | 1.288 (0.377) | 0.965 (0.209) | <.001, $\eta^2=0.658$ | .021 | .388 |
| | Rotation <sup>ab</sup> | 1.113 (0.238) | 1.704 (0.490) | 1.161 (0.207) | 1.189 (0.240) | 1.638 (0.434) | 1.271 (0.268) | <.001, $\eta^2=0.703$ | .347 | .130 |
| Left knee | FE <sup>a*</sup> | 1.859 (0.384) | 2.560 (0.669) | 1.966 (0.492) | 2.212 (0.393) | 2.735 (0.647) | 2.367 (0.657) | .001, $\eta^2=0.490$ | .001, $\eta^2=0.536$ | .328 |
| | VV <sup>a</sup> | 0.649 (0.101) | 0.883 (0.242) | 0.665 (0.080) | 0.693 (0.093) | 0.871 (0.205) | 0.729 (0.126) | .001, $\eta^2=0.537$ | .086 | .286 |
| Left ankle | DfPf <sup>a*</sup> | 1.518 (0.265) | 2.053 (0.359) | 1.540 (0.238) | 1.776 (0.323) | 2.192 (0.482) | 1.807 (0.432) | <.001, $\eta^2=0.619$ | .004, $\eta^2=0.465$ | .306 |
| | InvEv <sup>a</sup> | 1.072 (0.162) | 1.451 (0.272) | 1.106 (0.199) | 1.107 (0.168) | 1.400 (0.263) | 1.082 (0.198) | <.001, $\eta^2=0.730$ | .552 | .270 |
| Pelvis | Tilt <sup>a*</sup> | 0.788 (0.119) | 1.188 (0.407) | 0.835 (0.159) | 0.936 (0.178) | 1.242 (0.394) | 0.997 (0.157) | .006, $\eta^2=0.415$ | .001, $\eta^2=0.554$ | .225 |
| | Obliquity <sup>a</sup> | 0.778 (0.344) | 1.287 (0.911) | 0.826 (0.377) | 0.916 (0.454) | 1.269 (0.548) | 0.902 (0.352) | .001, $\eta^2=0.517$ | .109 | .444 |
| | Rotation <sup>a</sup> | 1.174 (0.316) | 2.064 (0.948) | 1.185 (0.287) | 1.292 (0.321) | 2.199 (1.247) | 1.301 (0.370) | <.001, $\eta^2=0.580$ | .183 | .997 |
| Trunk | FE <sup>a</sup> | 0.979 (0.250) | 1.417 (0.385) | 1.058 (0.274) | 1.142 (0.197) | 1.560 (0.683) | 1.131 (0.255) | .002, $\eta^2=0.469$ | .015 | .670 |
| | Bending <sup>a*</sup> | 0.834 (0.189) | 1.350 (0.383) | 0.787 (0.176) | 1.028 (0.299) | 1.537 (0.586) | 0.993 (0.302) | <.001, $\eta^2=0.580$ | .001, $\eta^2=0.647$ | .970 |
| | Rotation <sup>a</sup> | 1.339 (0.382) | 3.470 (2.612) | 1.352 (0.318) | 1.521 (0.350) | 3.494 (2.461) | 1.478 (0.404) | .001, $\eta^2=0.552$ | .646 | .866 |

FE: flexion/extension, AA: abduction/adduction, VV: valgus/varus, DfPf : dorsiflexion/plantarflexion, InvEv: inversion/eversion
<sup>a</sup> post-hoc difference between normal and active swing, <sup>b</sup> post-hoc difference between normal and held swing, \* symmetry difference

Significant effects of Swing were found on meanSD of all individual joint angles (p = 0.001-0.006, specific effect sizes in Table 1). Post-hoc tests revealed that, relative to normal swing, meanSD of each joint angle increased during active swing (p < 0.010) and that meanSD of left hip rotation increased when the arms were held (p = 0.009).

Significant effects of Symmetry were found on meanSD of trunk bending (p = 0.001, η^2^ = 0.647), pelvis tilt (p = 0.001, η^2^ = 0.554), right hip flexion (p < 0.001, η^2^ = 0.650), right hip abduction (p = 0.001, η^2^ = 0.529), right knee flexion (p < 0.001, η^2^ = 0.680), right knee valgus (p < 0.001, η^2^ = 0.601), right ankle dorsiflexion (p = 0.003, η^2^ = 0.469), left hip flexion (p = 0.008, η^2^ = 0.404), left knee flexion (p = 0.001, η^2^ = 0.536), and left ankle dorsiflexion (p = 0.004, η^2^ = 0.465). For all significant effects, meanSD increased in asymmetric gait relative to symmetric gait, indicating increased magnitude of variability with gait asymmetry.

### 3.2 λ_max_, local dynamic stability (Table 2)

There were no significant Swing^*^Symmetry interactions on λ_max_ (p ≥ 0.010).

**Table 2.** Maximum short-term finite-time Lyapunov exponents (λ_max_) of joint angles performed under six gait conditions varying by arm swing (normal, active, held) and symmetry (symmetric, asymmetric). Values are group means (inter-individual standard deviation). P-values of main and interaction effects are provided, with effect sizes (η^2^) for significant effects (p < 0.010).

| Joint | Degree of freedom | Symmetric |  |  | Asymmetric |  |  | Swing | Symmetry | Interaction |
| --- | --- | --- | --- | --- | --- | --- | --- | --- | --- | --- |
|  |  | Normal | Active | Held | Normal | Active | Held |  |  |  |
| Right hip | FE<sup>*</sup> | 2.501 (0.190) | 2.606 (0.220) | 2.579 (0.231) | 2.720 (0.202) | 2.752 (0.197) | 2.762 (0.209) | .142 | <.001, $\eta^2=0.835$ | .435 |
| | AA<sup>a*</sup> | 1.857 (0.107) | 1.975 (0.143) | 1.886 (0.128) | 1.946 (0.128) | 2.042 (0.150) | 1.983 (0.153) | .002, $\eta^2=0.424$ | <.001, $\eta^2=0.739$ | .782 |
|  | Rotation | 1.580 (0.158) | 1.627 (0.159) | 1.582 (0.135) | 1.643 (0.144) | 1.664 (0.166) | 1.652 (0.172) | .227 | .014 | .620 |
| Right knee | FE<sup>*</sup> | 2.667 (0.191) | 2.657 (0.207) | 2.716 (0.165) | 2.835 (0.164) | 2.748 (0.116) | 2.886 (0.206) | .007, $\eta^2=0.299$ | <.001, $\eta^2=0.713$ | .255 |
| | VV<sup>*</sup> | 1.576 (0.204) | 1.620 (0.227) | 1.575 (0.130) | 1.666 (0.200) | 1.623 (0.172) | 1.698 (0.193) | .852 | .002, $\eta^2=0.511$ | .017 |
| Right ankle | DfPf<sup>*</sup> | 2.095 (0.134) | 2.176 (0.128) | 2.113 (0.126) | 2.160 (0.101) | 2.165 (0.086) | 2.212 (0.121) | .217 | .009, $\eta^2=0.397$ | .047 |
| | InvEv<sup>a</sup> | 1.558 (0.158) | 1.692 (0.130) | 1.584 (0.153) | 1.611 (0.147) | 1.772 (0.142) | 1.611 (0.130) | <.001, $\eta^2=0.612$ | .022 | .276 |
| Left hip | FE<sup>*</sup> | 2.472 (0.166) | 2.604 (0.231) | 2.549 (0.211) | 2.639 (0.177) | 2.677 (0.200) | 2.673 (0.193) | .077 | <.001, $\eta^2=0.598$ | .298 |
|  | AA | 1.901 (0.187) | 1.987 (0.188) | 1.914 (0.187) | 1.957 (0.154) | 2.025 (0.184) | 1.987 (0.213) | .054 | .018 | .589 |
|  | Rotation | 1.556 (0.108) | 1.657 (0.162) | 1.562 (0.106) | 1.524 (0.108) | 1.605 (0.191) | 1.577 (0.119) | .026 | .195 | .046 |
| Left knee | FE | 2.669 (0.199) | 2.646 (0.205) | 2.738 (0.175) | 2.742 (0.178) | 2.685 (0.115) | 2.778 (0.190) | .005, $\eta^2=0.314$ | .085 | .817 |
|  | VV | 1.538 (0.199) | 1.541 (0.216) | 1.540 (0.183) | 1.523 (0.177) | 1.545 (0.180) | 1.594 (0.219) | .589 | .563 | .268 |
| Left ankle | DfPf<sup>*</sup> | 2.038 (0.134) | 2.096 (0.146) | 2.077 (0.132) | 2.107 (0.135) | 2.152 (0.154) | 2.157 (0.125) | .097 | .006, $\eta^2=0.425$ | .849 |
|  | InvEv | 1.620 (0.168) | 1.713 (0.126) | 1.642 (0.164) | 1.672 (0.178) | 1.726 (0.060) | 1.654 (0.156) | .025 | .274 | .395 |
| Pelvis | Tilt | 2.464 (0.127) | 2.466 (0.166) | 2.519 (0.201) | 2.500 (0.160) | 2.519 (0.206) | 2.566 (0.141) | .106 | .023 | .951 |
|  | Obliquity | 2.215 (0.186) | 2.288 (0.273) | 2.239 (0.217) | 2.260 (0.171) | 2.322 (0.200) | 2.291 (0.177) | .142 | .145 | .913 |
|  | Rotation | 2.348 (0.127) | 2.394 (0.159) | 2.360 (0.168) | 2.377 (0.129) | 2.450 (0.198) | 2.354 (0.203) | .112 | .112 | .684 |
| Trunk | FE | 2.715 (0.150) | 2.692 (0.162) | 2.720 (0.144) | 2.715 (0.143) | 2.797 (0.138) | 2.780 (0.125) | .540 | .023 | .041 |
|  | Bending | 2.941 (0.166) | 2.849 (0.208) | 2.904 (0.233) | 3.014 (0.210) | 2.893 (0.260) | 2.945 (0.171) | .107 | .030 | .842 |
|  | Rotation | 2.656 (0.145) | 2.573 (0.208) | 2.688 (0.181) | 2.742 (0.174) | 2.617 (0.221) | 2.686 (0.147) | .039 | .119 | .582 |

FE: flexion/extension, AA: abduction/adduction, VV: valgus/varus, DfPf : dorsiflexion/plantarflexion, InvEv: inversion/eversion
<sup>a</sup> post-hoc difference between normal and active swing, <sup>b</sup> post-hoc difference between normal and held swing, <sup>\*</sup> symmetry difference

Swing effects on λ_max_ were observed for right hip abduction (p = 0.002, η^2^ = 0.424) and right ankle inversion (p < 0.001, η^2^ = 0.612). Post-hoc comparisons revealed that, relative to normal swing, λ_max_ increased during active swing (p < 0.010) but did not change when the arms were held. Effects of Swing were also observed for right knee flexion (p = 0.007, η^2^ = 0.299) and left knee flexion (p = 0.005, η^2^ = 0.314), but these post-hoc comparisons with normal swing were not statistically significant (p ≥ 0.010).

Symmetry effects on λ_max_ were observed for right hip flexion (p < 0.001, η^2^ = 0.835), right hip abduction (p < 0.001, η^2^ = 0.739), right knee flexion (p < 0.001, η^2^ = 0.713), right knee valgus (p = 0.002, η^2^ = 0.511), right ankle dorsiflexion (p = 0.009, η^2^ = 0.397), left hip flexion (p < 0.001, η^2^ = 0.598), and left ankle dorsiflexion (p = 0.006, η^2^ = 0.425). In all cases, λ_max_ increased during asymmetric gait relative to symmetric gait, indicating decreased local dynamic stability with asymmetry. Trends for increased λ_max_ of trunk flexion (p = 0.023) and trunk bending (p = 0.030) with asymmetry were supported by a significant increase in λ_max_ of Euclidean norm trunk angle (p = 0.002, η^2^ = 0.502) (Supplementary Table 1).

### 3.3 DFAα, statistical persistence (Table 3)

There were no significant effects on DFAα of any joint angles (p ≥ 0.010). DFAα means in symmetric and asymmetric gait conditions were all > 0.500, with some means closer to 1.000 than to 0.500 (trunk: 0.532-0.805, pelvis: 0.568-0.717, hips: 0.576-0.840, knees: 0.633-0.844, ankles: 0.587-0.698), indicating that range of motion fluctuations were persistent on average.

**Table 3.** DFA scaling exponent of range of motion (DFAα) of joint angles performed under six gait conditions varying by arm swing (normal, active, held) and symmetry (symmetric asymmetric). Values are group means (inter-individual standard deviation). P-values of main and interaction effects are provided, with effect sizes (η^2^) for significant effects (p < 0.010).

| Joint | Degree of freedom | Symmetric |  |  | Asymmetric |  |  | Swing | Symmetry | Interaction |
| --- | --- | --- | --- | --- | --- | --- | --- | --- | --- | --- |
|  |  | Normal | Active | Held | Normal | Active | Held |  |  |  |
| Right hip | FE | 0.650 (0.174) | 0.822 (0.183) | 0.707 (0.223) | 0.744 (0.177) | 0.773 (0.227) | 0.708 (0.192) | .140 | .706 | .363 |
|  | AA | 0.644 (0.136) | 0.645 (0.148) | 0.723 (0.198) | 0.676 (0.167) | 0.652 (0.174) | 0.674 (0.206) | .272 | .911 | .701 |
|  | Rotation | 0.627 (0.120) | 0.621 (0.150) | 0.649 (0.150) | 0.635 (0.121) | 0.650 (0.180) | 0.586 (0.171) | .887 | .773 | .505 |
| Right knee | FE | 0.633 (0.162) | 0.725 (0.175) | 0.692 (0.097) | 0.721 (0.185) | 0.785 (0.166) | 0.766 (0.209) | .306 | .048 | .940 |
|  | VV | 0.670 (0.163) | 0.679 (0.189) | 0.758 (0.139) | 0.712 (0.162) | 0.675 (0.189) | 0.728 (0.173) | .393 | .951 | .541 |
| Right ankle | DfPf | 0.660 (0.194) | 0.664 (0.180) | 0.658 (0.185) | 0.658 (0.212) | 0.662 (0.213) | 0.664 (0.156) | .997 | .976 | .996 |
|  | InvEv | 0.668 (0.134) | 0.636 (0.133) | 0.698 (0.149) | 0.630 (0.175) | 0.587 (0.163) | 0.595 (0.182) | .444 | .133 | .699 |
| Left hip | FE | 0.719 (0.168) | 0.768 (0.255) | 0.740 (0.159) | 0.682 (0.187) | 0.840 (0.237) | 0.710 (0.199) | .215 | .959 | .497 |
|  | AA | 0.645 (0.183) | 0.630 (0.208) | 0.717 (0.211) | 0.669 (0.212) | 0.688 (0.173) | 0.661 (0.213) | .813 | .816 | .534 |
|  | Rotation | 0.576 (0.153) | 0.549 (0.202) | 0.598 (0.196) | 0.629 (0.121) | 0.687 (0.183) | 0.607 (0.121) | .935 | .076 | .206 |
| Left knee | FE | 0.734 (0.209) | 0.748 (0.159) | 0.651 (0.163) | 0.703 (0.257) | 0.844 (0.205) | 0.776 (0.262) | .340 | .074 | .339 |
|  | VV | 0.711 (0.193) | 0.669 (0.182) | 0.715 (0.205) | 0.782 (0.136) | 0.712 (0.251) | 0.699 (0.177) | .517 | .256 | .722 |
| Left ankle | DfPf | 0.693 (0.171) | 0.683 (0.186) | 0.641 (0.163) | 0.637 (0.203) | 0.599 (0.160) | 0.688 (0.204) | .827 | .427 | .230 |
|  | InvEv | 0.633 (0.169) | 0.653 (0.224) | 0.632 (0.204) | 0.618 (0.161) | 0.615 (0.156) | 0.693 (0.169) | .781 | .935 | .377 |
| Pelvis | Tilt | 0.568 (0.140) | 0.714 (0.159) | 0.633 (0.229) | 0.585 (0.149) | 0.618 (0.156) | 0.606 (0.175) | .259 | .324 | .170 |
|  | Obliquity | 0.611 (0.173) | 0.663 (0.172) | 0.697 (0.227) | 0.675 (0.246) | 0.664 (0.241) | 0.616 (0.219) | .946 | .894 | .364 |
|  | Rotation | 0.672 (0.149) | 0.710 (0.204) | 0.749 (0.157) | 0.646 (0.174) | 0.717 (0.184) | 0.691 (0.139) | .404 | .410 | .721 |
| Trunk | FE | 0.532 (0.080) | 0.623 (0.209) | 0.574 (0.104) | 0.616 (0.159) | 0.621 (0.149) | 0.674 (0.133) | .408 | .043 | .174 |
|  | Bending | 0.616 (0.098) | 0.646 (0.136) | 0.684 (0.198) | 0.650 (0.233) | 0.664 (0.206) | 0.605 (0.144) | .772 | .835 | .300 |
|  | Rotation | 0.676 (0.121) | 0.805 (0.197) | 0.696 (0.209) | 0.617 (0.211) | 0.694 (0.170) | 0.623 (0.148) | .062 | .018 | .872 |

DFA: detrended fluctuation analysis, FE: flexion/extension, AA: abduction/adduction, VV: valgus/varus, DfPf : dorsiflexion/plantarflexion, InvEv: inversion/eversion
<sup>a</sup> post-hoc difference between normal and active swing, <sup>b</sup> post-hoc difference between normal and held swing, \* symmetry difference

### 3.4 SaEn, regularity (Table 4)

There were no significant Swing^*^Symmetry interactions on SaEn (p ≥ 0.010).

**Table 4.**
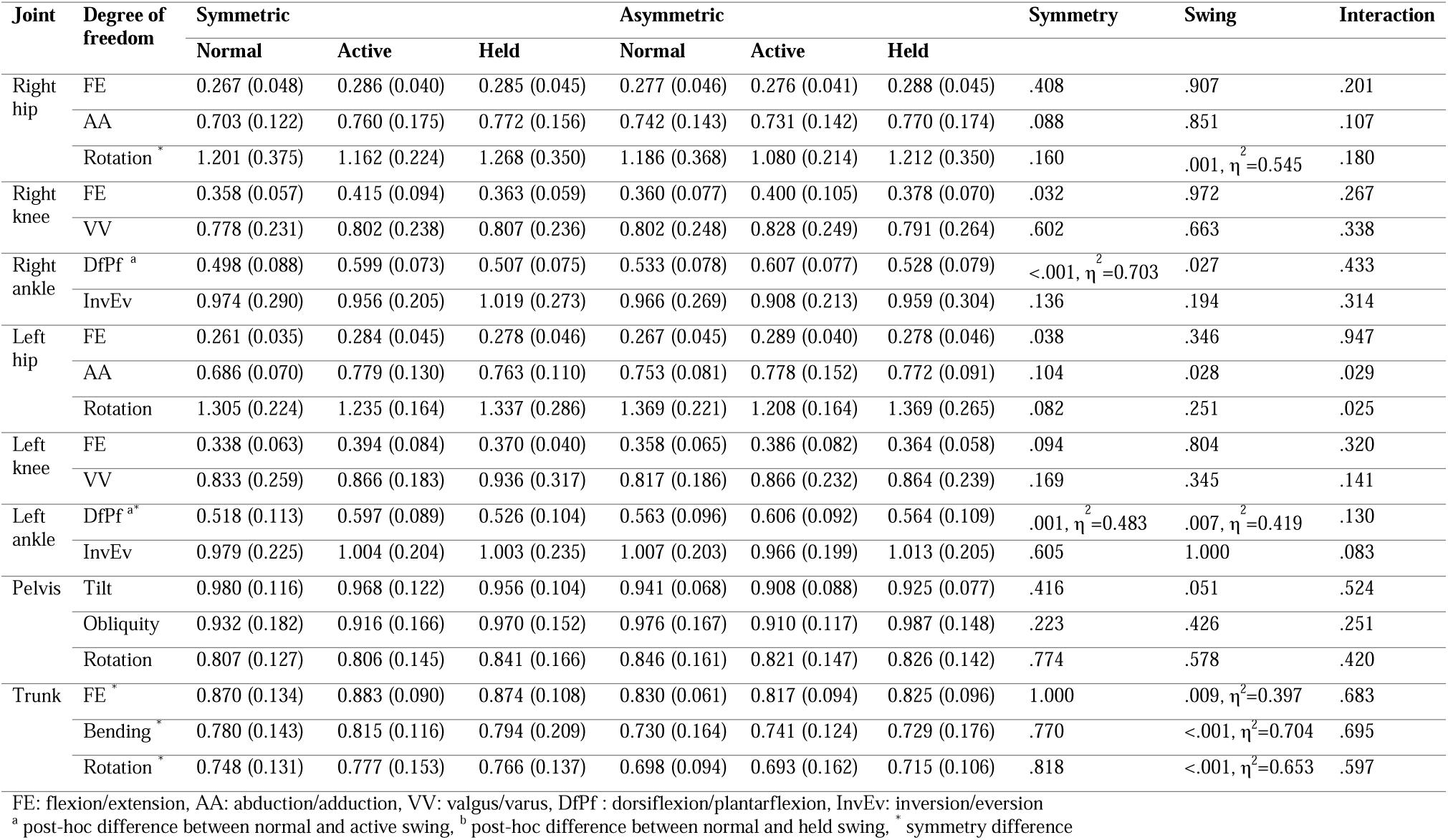
Sample entropy (SaEn) of joint angles performed under six gait conditions varying by arm swing (normal, active, held) and symmetry (symmetric, asymmetric). Values are group means (inter-individual standard deviation). P-values of main and interaction effects are provided, with effect sizes (η^2^) for significant effects (p < 0.010).

Significant Swing effects showed that, compared to normal swing, active swing led to increased SaEn of right ankle dorsiflexion (p < 0.001, η^2^ = 0.703) and left ankle dorsiflexion (p = 0.007, η^2^ = 0.419). No significant changes in SaEn were observed when the arm was held (p ≥ 0.010).

Significant Symmetry effects showed that, compared to symmetric gait, asymmetric gait led to decreased SaEn of trunk flexion (p = 0.009, η^2^ = 0.397), trunk bending (p < 0.001, η^2^ = 0.704), trunk rotation (p < 0.001, η^2^ = 0.653), and right hip rotation (p = 0.001, η^2^ = 0.545), and increased SaEn of left ankle dorsiflexion (p = 0.007, η^2^ = 0.419).

All significant effects reported in this study had a large effect size (η^2^ ≥ 0.14).

## 4. Discussion

### 4.1 Active arm swing altered variability of lower limb joint angles while preserving pelvis and trunk stability and regularity

In contrast with our hypothesis and the findings of Wu et al. (2016), active arm swing increased the magnitude of variability in tridimensional hip, knee, and ankle joint angles. This is more in line with the increased magnitude of variability in step time, step length, and step width reported in previous investigations of our sample (Hill and Nantel, 2019; Siragy et al., 2020). However, unlike in Hill & Nantel (2019) where local dynamic stability of trunk linear and angular velocities decreased with active arm swing, local dynamic stability of trunk angles did not significantly change. This difference may be attributed to the different state-space constructions (linear and angular velocities together vs. uniaxial angles) for calculation of the maximum finite-time Lyapunov exponent (Gates and Dingwell, 2009). There was a trend for trunk rotation angle (p = 0.039) that was not significant after adjusting critical alpha for false discovery rate, so it is possible that a small or medium effect size may exist but went undetected. Yet, our findings and those of Hill and Nantel (2019) agree that trunk and pelvis stability are, at a minimum, preserved during active arm swing gait. The preservation of trunk and pelvis stability may be related to the more variable and dynamic base of support (Hill and Nantel, 2019), such that variability in the base of support would be dictated by variability in lower limb joint movements. In fact, local dynamic stability of the hip and ankle kinematics and regularity of the ankle kinematics decreased with active arm swing, indicating key alterations in the non-linear dynamics of lower limb joint movements; these may reflect the use of redundant movement patterns to preserve local trunk and pelvis stability during active arm swing.

Interestingly, influences of active arm swing on stride-to-stride lower limb dynamics were plane-specific, where joint angle stability decreased in the frontal plane (right hip abduction and ankle inversion) while regularity decreased only in the sagittal plane (right and left ankle dorsiflexion). This suggests that increased arm swing amplitude leads to stride-to-stride adjustments that preserve dynamic stability of joint movements in the main plane of motion. Plane-dependent adjustments have also been seen in the magnitude of ankle angle variability as a function of old age, with lower variability in the sagittal plane but higher variability in the frontal plane with older age (Bailey et al., 2020). Arm swing amplitude may therefore have a role in the plane-dependent stride-to-stride control patterns emerging with old age.

Restricted (held) arm swing had little to no effect on variability of lower limb joint angles, in agreement with the lack of effect seen on the magnitude of variability and the local dynamic stability of the trunk seen previously in steady-state gait (Bruijn et al., 2010; Hill and Nantel, 2019; Siragy et al., 2020). The only effect seen was an increase in the magnitude of variability of the left hip rotation. However, since this was not observed in the right hip and no other motor variability features were affected, this should be investigated in future studies by directly comparing the left and right sides. Restricted arm swing has been similarly found to have little effect on average gait kinematics, where Umberger (2008) reported similar hip, knee, and ankle joint angles in a gait cycle compared to regular arm swing (5.5-10.2% root mean square difference). As noted in that study, restricted arm swing was associated with different magnitude and shape of the free vertical moment at the foot, and of knee joint moment and power, suggesting that restricted arm swing has a greater influence on lower limb kinetics than kinematics during gait.

### 4.2 Arm swing amplitude did not influence asymmetry-related changes in lower limb variability patterns

In agreement with our second hypothesis, we found no significant interactions between arm swing amplitude and lower limb asymmetry on motor variability of trunk, pelvis, and lower limb joint angles. Independent of arm swing amplitude, lower limb asymmetry had a significant influence on motor variability of trunk, pelvis, and lower limb joint angles. Similar to trunk velocity (Hill and Nantel, 2019; Siragy et al., 2020), magnitude of variability increased for all joint angles and local dynamic stability decreased for Euclidean norm trunk angle and all lower limb joint angles. These lower limb adjustments were seen predominantly in sagittal plane degrees of freedom bilaterally. There was also statistical persistence in the range of motion fluctuations, and the regularity of right ankle dorsiflexion/plantarflexion decreased while regularity of all trunk angles and of the right hip rotation increased. Collectively, these findings suggest that individuals continually searched for motor strategies amongst their redundancy to perform asymmetric gait, particularly in the sagittal plane. Split-belt asymmetry adds a cognitive demand that, in agreement with McFadyen et al. (McFadyen et al., 2009), individuals seem to adjust to by coordinating gait timing through conscious attention to the redundancies of the lower extremity. Our findings provide two new insights into where this attention is specifically directed in the lower limbs. First, we saw more prevalent changes in magnitude of variability, local dynamic stability, and regularity in the right lower extremity (moving at the slower 0.96 m/s speed) suggesting that the attention to exploiting kinematic redundancies is more directed to the limb undergoing the change in speed; however, this study was not powered to also explore right-left differences and this needs to be confirmed statistically in future work. Second, we saw some changes at the hip and ankle that were not observed at the knee, suggesting that attention is directed more to joints with larger kinematic redundancies. Of note, these observations were at a lower asymmetry ratio (0.8:1) relative to the 1:2 ratio often employed in split-belt gait assessments (Hirata et al., 2019; McFadyen et al., 2009), indicating that even minor asymmetries can alter lower limb motor variability in healthy young adults. Our findings could be amplified in older adults, as healthy males and females aged 73.4 ± 4.7 years adapted less and more slowly to asymmetric gait induced by a split-belt treadmill than young adults (Bruijn et al., 2012). With healthy older age also associated with changes in gait asymmetry overground (Aboutorabi et al., 2016) and altered magnitude of variability in lower limb joint angles and muscle activation amplitude (Bailey et al., 2019, 2020), asymmetry may be a potential mechanism contributing to age-related adjustments in motor variability during gait.

### 4.3 Limitations

Findings are from young adults walking on a treadmill at 1.2 m/s; influences of arm swing and asymmetry on joint angle variability patterns may differ for older adults and different gait conditions. For instance, the treadmill gait produces lower variability of stride time (Hollman et al., 2016) and joint angles (Dingwell et al., 2001) relative to overground gait, which could mean that the reported adjustments related to arm swing and lower limb were also smaller in magnitude than those occurring overground. Since arm swing amplitude was manipulated by artificial conditions that maintained swing cadence but decreased interlimb coordination (Hill and Nantel, 2019), future investigations may wish to explore effects of more natural variations in arm swing amplitude between individuals using a correlation approach. Finally, males and females were grouped together for sufficient statistical power; given evidence of some differences in motor variability patterns of males and females during gait (Bailey et al., 2019, 2020), further work is needed to examine how sex influences arm swing and asymmetry-related adjustments in motor variability.

### 4.4 Conclusion

Active arm swing increased the magnitude of variability in joint angles across all planes, and, while preserving local dynamic stability and regularity of trunk and pelvis angles, decreased lower limb joint angle local dynamic stability in the frontal and transverse planes and regularity in the sagittal plane. Lower limb asymmetry increased the magnitude of variability of all joint angles and decreased the local dynamic stability of Euclidean norm trunk angle and lower limb joint angles, and decreased the regularity of right ankle dorsiflexion/plantarflexion while increasing regularity of all trunk angles. We conclude that young adults actively search for motor strategies amongst the redundancies of their ankle, knee, hip when actively swinging their arm and when walking asymmetrically, preserving stability and regularity of the pelvis and trunk during active arm swing but not during lower limb asymmetry. Findings may help explain arm swing amplitude and motor variability adjustments observed in gait of older adults but require confirmation in this population.

## Acknowledgement

The authors gratefully acknowledge Courtney Bridgewater for her assistance with data collection and the participants for their time.

---

**Supplementary Table 1.**
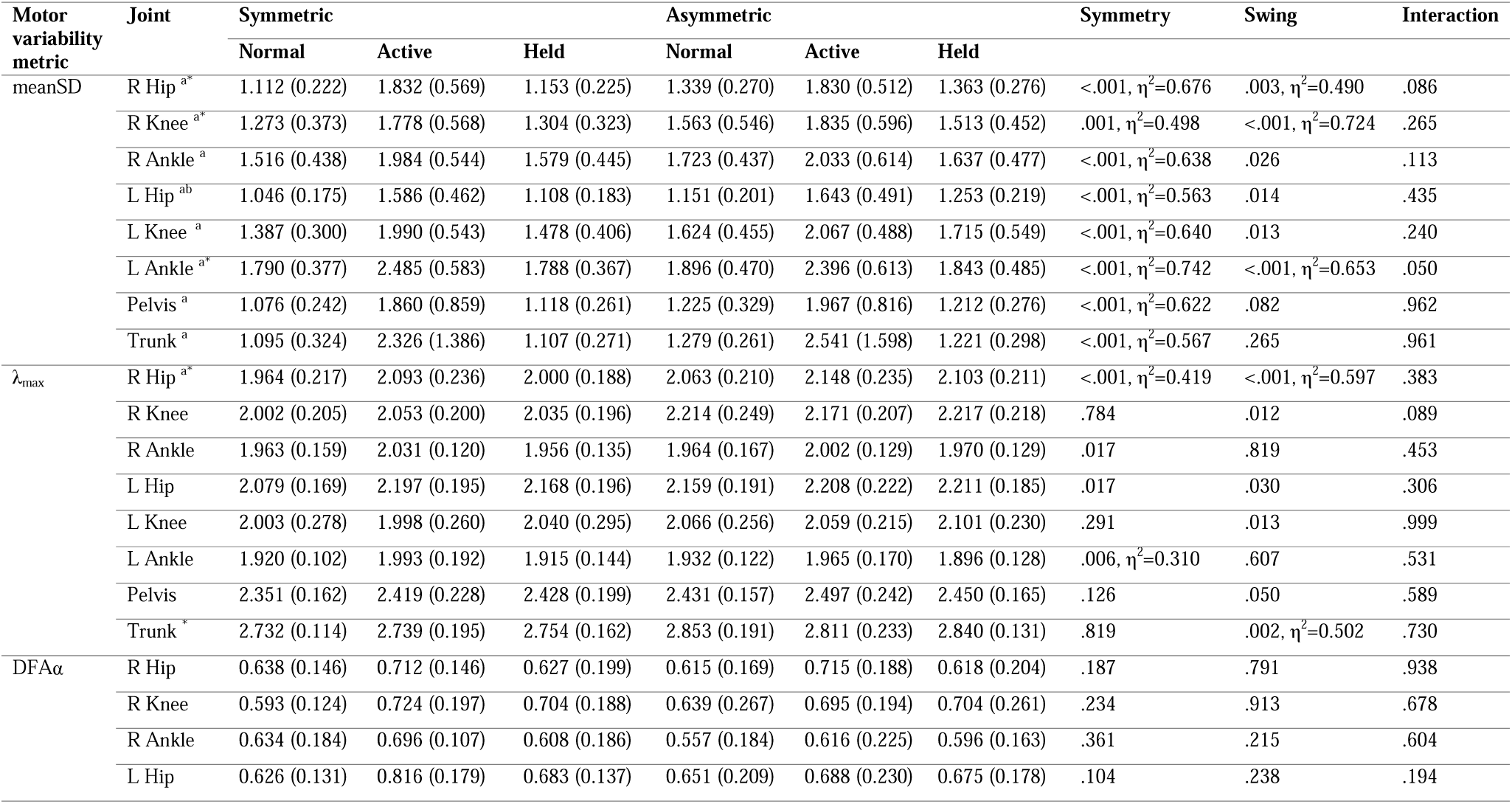
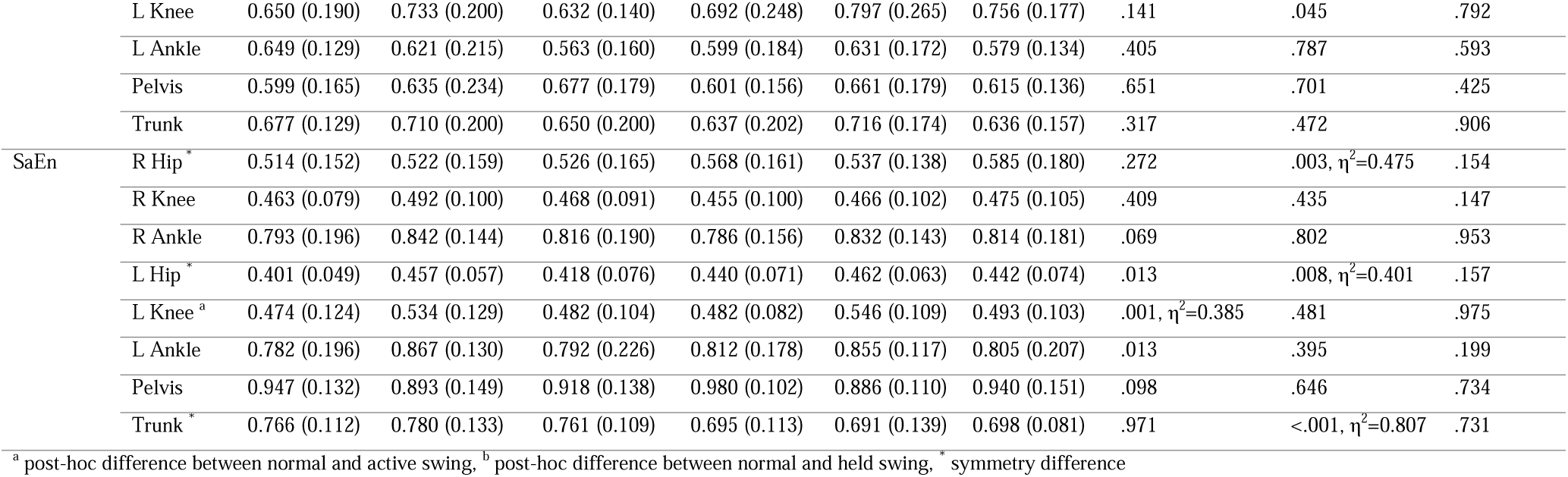
Motor variability of Euclidian mean joint angles in six gait conditions varying by arm swing (normal, active, held) and symmetry (symmetric, asymmetric). Metrics are mean standard deviation (meanSD), maximum short-term finite-time Lyapunov exponent (λ_max_), detrended fluctuation analysis scaling exponent of range of motion (DFAα), and sample entropy (SaEn). Values are group means (inter-individual standard deviation). P-values of main and interaction effects are provided, with effect sizes (η^2^) for significant effects (p < 0.010).

